# DGRNA: a long-context RNA foundation model with bidirectional attention Mamba2

**DOI:** 10.1101/2024.10.31.621427

**Authors:** Ye Yuan, Qushuo Chen, Xiaoyong Pan

**Affiliations:** Key Laboratory of Biopharmaceutical Preparation and Delivery, Chinese Academy of Sciences, Beijing, China; DigitalGene, Ltd, Shanghai, 200240, China; Institute of Image Processing and Pattern Recognition, Shanghai Jiao Tong University, and Key Laboratory of System Control and Information Processing, Ministry of Education of China, Shanghai, 200240, China

## Abstract

Ribonucleic acid (RNA) is an important biomolecule with diverse functions i.e. genetic information transfer, regulation of gene expression and cellular functions. In recent years, the rapid development of sequencing technology has significantly enhanced our understanding of RNA biology and advanced RNA-based therapies, resulting in a huge volume of RNA data. Data-driven methods, particularly unsupervised large language models, have been used to automatically hidden semantic information from these RNA data. Current RNA large language models are primarily based on Transformer architecture, which cannot efficiently process long RNA sequences, while the Mamba architecture can effectively alleviate the quadratic complexity associated with Transformers. In this study, we propose a large foundational model DGRNA based on the bidirectional Mamba trained on 100 million RNA sequences, which has demonstrated exceptional performance across six RNA downstream tasks compared to existing RNA language models.

## Introduction

Ribonucleic acids (RNAs) are fundamental macromolecules with versatile roles beyond their traditional function as intermediaries in genetic information transfer[1, 2]. Within the framework of the central dogma, RNA is categorized into three primary types: messenger RNA (mRNA), which carries genetic information from DNA to ribosomes for protein synthesis; ribosomal RNA (rRNA), which constitutes the core of ribosomes and facilitates protein synthesis; and transfer RNA (tRNA), which translates mRNA sequences into amino acids. Additionally, RNAs can be divided into protein-coding and non-coding RNAs[3-5]. Non-coding RNAs (ncRNAs), such as microRNAs (miRNAs) and long non-coding RNAs (lncRNAs), do not encode proteins but instead regulate gene expression and contribute to various cellular functions. miRNAs are involved in regulating gene expression post-transcriptionally, while lncRNAs participate in processes like chromatin remodeling and epigenetic regulation[6-9]. knowledge and RNA-based therapeutic applications. [10-12].

The rise of high-throughput sequencing technologies has generated vast amounts of unlabeled RNA sequence data, creating opportunities for the development of sophisticated computational models. Meanwhile, the emergence of various databases has significantly propelled the application of numerous RNA-related algorithms. These databases, i.e. the MARS database, provide rich and diverse datasets that leverage to develop and refine computational methods for RNA analysis. Due to the limited number of RNA 3D structures and functions, unsupervised large RNA language models can be developed to predict RNA structure and function more effectively[13-16].

Currently, RNA language models are predominantly based on BERT, which is a pre-trained language model utilizing the Transformer architecture to understand contextual relationships. By employing a bidirectional encoder, BERT simultaneously considers information from both sides of a word, enhancing its language comprehension capabilities[17-19]. RNA-FM, as the first pre-trained model in the RNA domain, marks a significant advancement in RNA sequence analysis by leveraging self-supervised learning on an extensive dataset of 23 million non-coding RNA sequences, RNA-FM’s architecture, which consists of 12 Transformer encoder blocks, enables to capture intricate relationships within RNA sequences, making it particularly effective for downstream tasks such as secondary and tertiary structure prediction. In the fine-tuning phase, RNA-FM is applied to various downstream tasks, including SARS-CoV-2 genome analysis, protein-RNA interaction prediction, and functional prediction of mRNA untranslated regions. RNA-FM demonstrates exceptional performance in these tasks, for instance, in RNA secondary structure prediction, RNA-FM achieves up to a 30% improvement in F1 scores compared to the state-of-the-art methods[20]. Since the release of the RNA foundational model RNA-FM in 2020, several large RNA language models have been developed, significantly enhancing our ability to analyze and predict RNA structures and functions. One notable model is RNABERT, which adapts the BERT architecture specifically for RNA sequences, allowing for better context-aware embeddings. This model excels in tasks like sequence classification and provides insights into RNA’s functional roles by capturing contextual relationships within RNA sequences[21]. Another important model is RNA-MSM, which employs multiple sequence alignments (MSA) to capture evolutionary information, making it particularly effective for tasks like base pairing and secondary structure prediction. Its focus on evolutionary conservation to identify functional regions within RNAs[22]. Shortly after that, RNAErnie is developed as a novel pre-trained RNA language model based on the ERNIE framework, utilizing two key strategies to enhance its performance. First, it integrates RNA motifs as biological prior knowledge, applying random masking at the motif level and appending RNA type information (e.g., miRNA, lncRNA) to sequences during pre-training. Second, it introduces a type-guided fine-tuning strategy to address out-of-distribution tasks by predicting RNA types and appending them as auxiliary information[23]. As one of the largest RNA language models to date, RiNALMo is an RNA language model with 650 million parameters. It has been pre-trained on a vast array of non-coding RNA sequences. RiNALMo demonstrates remarkable generalization capabilities for RNA families not seen in the training data. The model leverages advanced Transformer architecture and modern techniques, such as rotary position encoding and the SwiGLU activation function, to enhance the efficiency of processing large-scale biological data. As a model trained on the large dataset with the broadest variety of RNA types[24], UNI-RNA, employs advanced techniques such as rotary embeddings and Flash Attention to enhance its processing of RNA sequences. This model achieves promising results in predicting the structures and functions of various RNA types, especially in cross-species splice site prediction[25].

However, the Transformer model, while excelling in handling sequence data due to its self-attention mechanism, suffers to quadratic growth in computational complexity as sequence length increases, leading to significant resource consumption for long RNA sequences. To address this challenge, Mamba employs global receptive fields and dynamic weighting strategies to effectively capture important information in long sequences while avoiding the quadratic computational complexity. This design allows Mamba to maintain model expressiveness while significantly reducing resource consumption. Not long after, Mamba 2 is introduced to further improve upon traditional Transformer models by incorporating the State Space Dual (SSD) framework. This framework reveals a profound connection between State Space Models (SSMs) and semi-separable matrices, which possess unique structural properties that enable efficient representation and processing of sequence data. In Mamba 2, the core layer enhancement is based on the selective SSM from the original Mamba model, resulting in a significant increase in training efficiency, achieving 2 to 8 times faster than its predecessor[26-28].

Considering the characteristics of long RNA sequences, we design a RNA foundation model DGRNA based on a bidirectional Mamba2 model on 100 million RNA sequences, achieving state-of-the-art (SOTA) performance across several downstream tasks. This extensive training allows the model DGRNA to effectively leverage the rich contextual information hidden in RNA sequences. DGRNA nont only outperforms existing RNA language models with 100 million parameters, but also demonstrates improved efficiency in processing long RNA sequences. These results highlight the model’s ability to integrate semantic information and biological insights hidden in RNA sequences, paving the way for advancements in RNA biology and potential RNA therapeutic applications.

## Results

### Overview of DGRNA

To handle long RNA sequences, we develop a pre-trained RNA language model called DGRNA based on Bidirectional Mamba. As shown in **Figure 1**, The model, built upon the Mamba 2 framework, consists of 12 bidirectional Mamba blocks and is followed by an attention mechanism at the end. DGRNA outputs an embedding with a dimension of 768. Pre-training is conducted on approximately processed 100 million RNA sequences derived from MARS database. Upon completion of the pre-training phase, we test the model across six downstream tasks with a total of eight benchmark datasets. These tasks and datasets allowed us to evaluate the model’s performance and generalization capabilities across various RNA-related analyses, i.e., RNA-RNA interaction prediction, RNA-protein binding site identification, and analysis of non-coding RNA functions. The results from these evaluations not only demonstrated the robustness of our pre-trained model DGRNA, but also highlighted its potential applications in advancing RNA research and therapeutic development. In downstream tasks, we employed a fine-tuning strategy, which is a common method for applying pre-trained models to specific tasks. This method leverages the general knowledge acquired during the pre-training phase, allowing the model to learn the nuances of the new task more effectively. Depending on the complexity of the downstream task and the amount of available data, certain layers of the model can be frozen or unfrozen.

**Figure 1:**
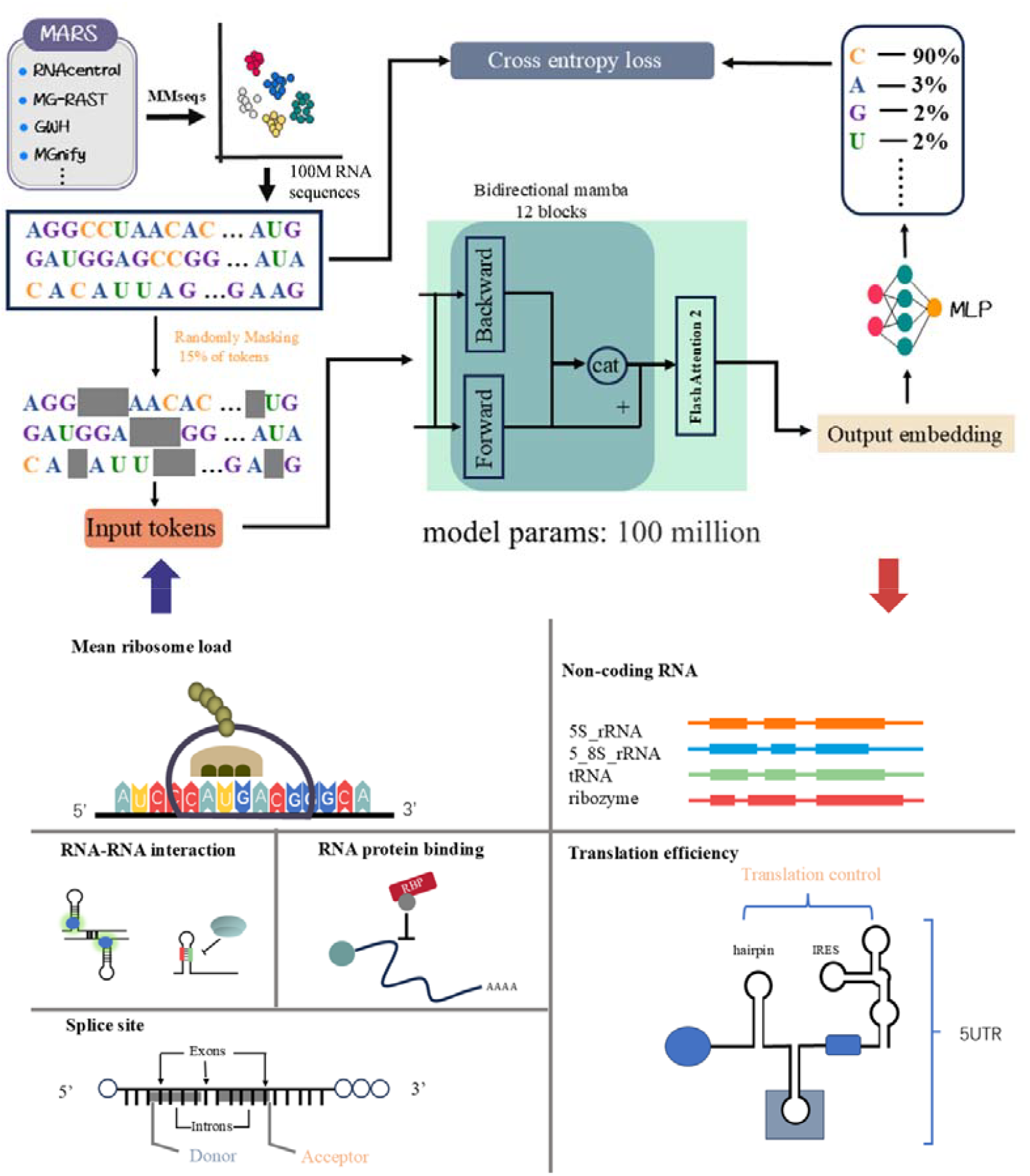
Overview of DGRNA pre-training and RNA-related downstream tasks. In the initial phase, DGRNA is trained on 100 million unlabeled RNA sequences sourced from MARS databases. This language model consists of 12 bidirectional Mamba 2 blocks with an attention mechanism, and has an output embedding dimension of 768. The number of the model’s parameters is 100 million (100M). DGRNA is evaluated for six RNA-related downstream tasks including mean ribosome load, non-coding RNA, RNA-RNA interaction, RNA protein binding, splice site and translation efficiency.

**Figure 2.**
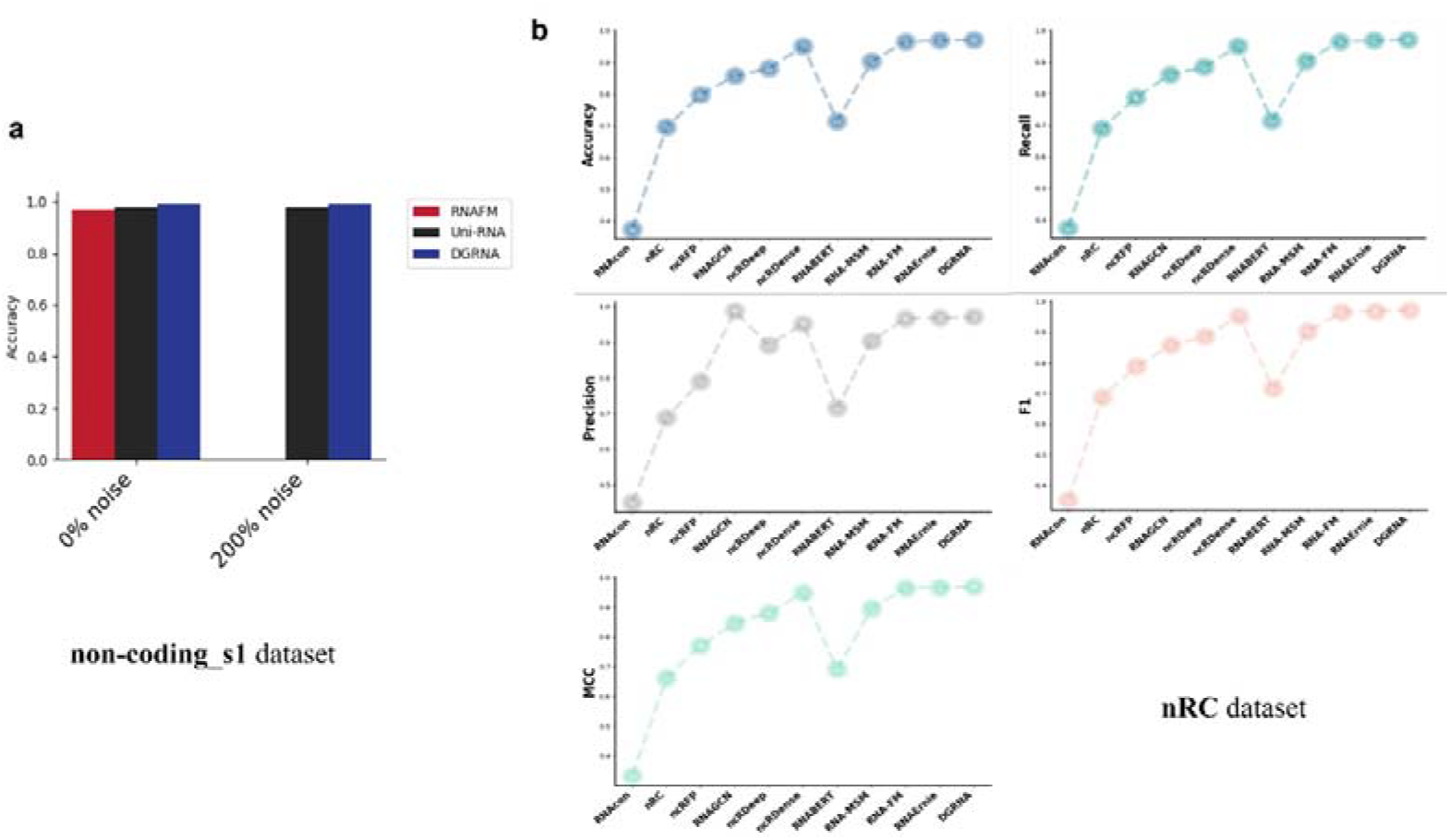
Performance of DGRNA and other RNA language models on the non-coding RNA classification task. a) The accuracy on two subsets of non-coding_s1 dataset; b) The performance of five metrics on nRC dataset

### Comparing DGRNA with other language models on Non-coding classification

The analysis of non-coding RNA provides insights into regulatory elements that govern gene expression. We utilized two well-established non-coding datasets to evaluate our method DGRNA: non-coding_s1 and nRC. In the non-coding_s1 task, we implement two linear layers combined with batch normalization layer to boost the model’s performance and stability. In the fine-tuning phase, we adjust the model parameters on a specific dataset, enabling it to adapt to specific task. This process involves freezing some layers while updating others, which helps retain learned features while allowing the model to refine its performance. As shown Figure 1a and Supplementary Table 1, we observe that the performance of DGRNA aligns closely with that of Uni-RNA in the non-coding_s1 task. The F1 score for both models reached 0.98, indicating an excellent balance between precision and recall. This high score underscores their strong performance in the task and suggests that both models are effective in feature extraction for ncRNA classification. In the nRC task, we also observed that our method DGRNA achieved the best performance in supplementary Table 2 and Figure 1b, outperforming existing RNA language models.

### The performance of DGRNA on 5UTR regression task

Following the RiNALMo approach, the outputs from the language model are processed through a subsequential layer comprising two ResNet convolutional blocks. Mean squared error is used as the loss function, and Adam serves as the optimizer with a total of 50 epochs for training. The evaluation metric used was R2, with a value close to 1 indicating a strong correlation. As shown in the **Table 1**, we found that the performance of DGRNA was consistent with RiNALMo, achieving an R2 score of 0.93. However, RiNALMo has a larger number of parameters, leading to slower training. This highlights the effectiveness of DGRNA in comparison to established language models.

**Table 1.** The different RNA foundation models evaluated on 5UTR regression task with correlation coefficient R2.

| Method | Random7600 | Human7600 |
| --- | --- | --- |
| RNA-FM | 0.85 | 0.79 |
| Uni-RNA | 0.91 | 0.85 |
| RiNALMo | **0.93** | 0.86 |
| DGRNA | **0.93** | **0.87** |

### DGRNA outperforms existing methods on RNA interaction prediction

Accurate predictions of RNA interactions can provide insights into regulatory mechanisms and enhance our understanding of biological processes like gene expression and translation. In this study, we evaluate DGRNA on two RNA interaction prediction tasks: RNA-RNA interaction and RNA-protein interaction. For RNA-RNA interaction, we use the key benchmark dataset DeepMirTar and employ the same methodology as RNAErnie to evaluate DGRNA’s performance. The goal of this task is to estimate interactions between RNA sequences, such as microRNA (miRNA) and messenger RNA (mRNA), and involves mapping two RNA sequences to binary labels (0 for no interaction and 1 for interaction)..Our approach utilizes the TBTH architecture for fine-tuning, similar to RNAErnie[29], integrating multiple components: a convolutional neural network (CNN), a bidirectional long short-term memory (Bi-LSTM) network, and a multi-layer perceptron (MLP). Small modifications include adjusting the learning rate and batch size. The results indicate that our model DGRNA outperforms all other methods as presented in **Supplementary Table 3** and **Figure 3a**. For the F1 metric, our model DGRNA achieved a 1% improvement than state-of-the-art methods, and it also demonstrated outstanding performance in accuracy, precision and AUC. The results underscore the DGRNA model’s potential for accurately predicting RNA-RNA interactions.

**Figure 3.**
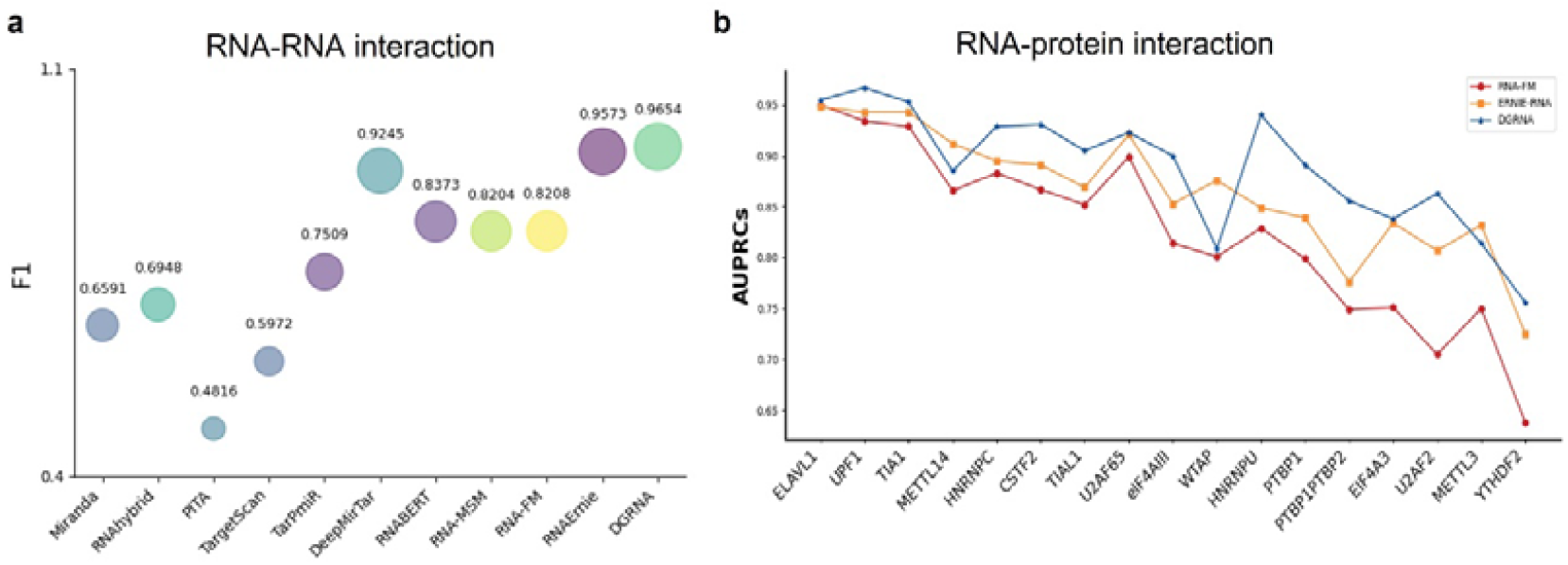
The performance of DGRNA on RNA interaction prediction. a) F1 score of various models on the RNA-RNA interaction dataset; b) AUCs of various language models on the RBP-RNA interaction dataset.

In addition, we adopted the data processing procedure of PrismNet, replacing the original network architecture with a two-layer MLP and substituting the icSHAPE input with the average of the embeddings, specifically averaging along the length dimension. During the fine-tuning phase, we initialized the learning rate to 1e-3 and employed the cross-entropy loss function to update the model parameters. Additionally, we set the tolerable number of training epochs to 20 for early stopping with a total of 200 epochs. As shown in the **Supplementary Table 4** and **Figure 3b**, we found that our method DGRNA achieved the best performance in 14 out of 17 RBP datasets and the average AUPRC (Area Under Precision-Recall Curve) across 17 RBPs is 0.889, which is higher than 0.824 of RNA-FM and 0.865 of ERNIE-RNA. Additionally, we observed an interesting phenomenon that the remaining three datasets, which performed poorly, all shared a common characteristic of having fewer than 2000 training samples.

### DGRNA is superior to other language models on Translation efficiency prediction task

Mean ribosome load prediction allows us to understand the translation efficiency of RNAs at the molecular level. In this study, we validate the performance using a 10-fold cross-validation approach. After obtaining the embeddings for the RNA sequences, we average them along the length dimension to create a fixed-length representation for each sequence, effectively summarizing the information contained in the embeddings while maintaining important features. We compare DGRNA with other RNA language models, such as RiNALMo, ERNIE-RNA, RNAERNIE[30], and RNABERT. As shown in **Table 5**, RiNALMo demonstrated the second-best performance in our analysis. Our method DGRNA achieved the best results with a Spearman correlation of 0.78, surpassing 0.74 of RiNALMo by 3%. This highlights the effectiveness of DGRNA in capturing the hidden information for RNA translation efficiency. Furthermore, our method significantly outperformed the last-place model, with a difference of nearly 40%. This substantial gap underscores the robustness and reliability of DGRNA in comparison to other RNA language models.

**Table 5.**
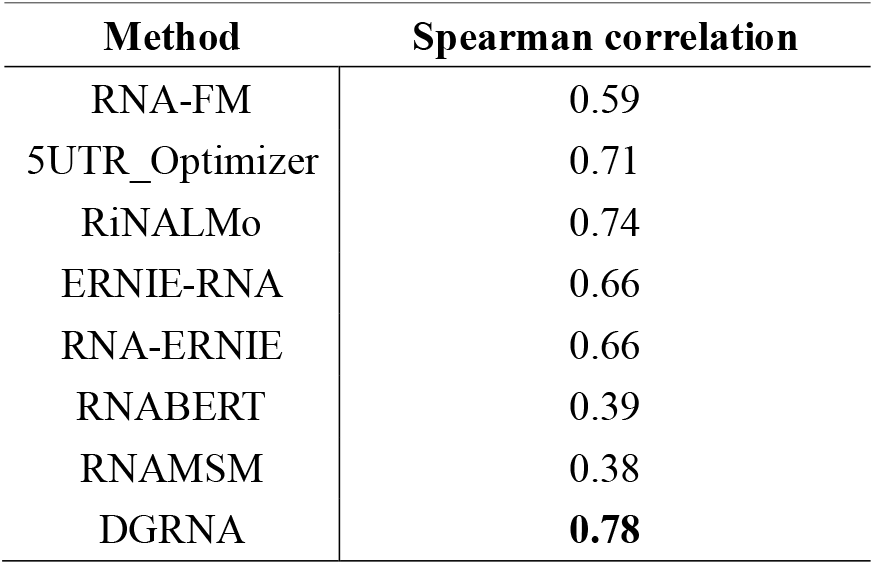
The different RNA foundation models were evaluated on the translation efficiency prediction task with Spearman correlation.

### Evaluating DGRNA on the Splicing Site Prediction task

Splice site identification is key to understanding alternative splicing, which has profound implications associated with protein diversity. To validate the reliability and generalizability of our model DGRNA, we employed SpliceBERT’s 10-fold cross-validation approach. We then conducted tests on four separate datasets encompassing zebrafish, fruit fly, worm, and Arabidopsis. We use the output of the “[CLS]” token from the last layer of DGRNA as input, passing it to a two-layer fully connected neural network for model fine-tuning. The ten models from 10-fold cross-validation are tested on independent dataset and then averaged. As shown in **Table 6**, we observed that on the Arabidopsis Acceptor dataset, DGRNA performed slightly lower than SpliceBERT, but on all other independent datasets, DGRNA achieved the best performance. The results also demonstrate the effectiveness of DGRNA on learning semantic RNA information from unlabeled RNA sequences.

**Table 6.** The different RNA foundation models were evaluated on Splicing Site prediction task with metric F1.

| Species | Type | RNA-FM | SpliceBERT | DGRNA |
| --- | --- | --- | --- | --- |
| zebrafish | Donor | 0.9538 | 0.9692 | **0.9780** |
|  | Acceptor | 0.9202 | 0.9443 | **0.9641** |
|  | Average | 0.9370 | 0.9568 | **0.9711** |
| fruit fly | Donor | 0.9385 | 0.9549 | **0.9657** |
|  | Acceptor | 0.8970 | 0.9373 | **0.9436** |
|  | Average | 0.9178 | 0.9461 | **0.9547** |
| worm | Donor | 0.9166 | 0.9411 | **0.9632** |
|  | Acceptor | 0.8735 | 0.9274 | **0.9321** |
|  | Average | 0.8950 | 0.9343 | **0.9477** |
| arabidopsis | Donor | 0.9075 | 0.9380 | **0.9677** |
|  | Acceptor | 0.8694 | **0.9343** | 0.9337 |
|  | Average | 0.8884 | 0.9361 | **0.9507** |

## Conclusion

In this study, we introduce DGRNA, a pre-trained RNA model based on the Mamba framework, which has demonstrated promising potential in addressing a variety of downstream tasks crucial to RNA biology. By leveraging the MARS dataset, which comprises 100 million RNA sequences, DGRNA is trained on these RNA sequences to capture intricate RNA features and structural information. DGRNA’s performance on these downstream tasks achieves state-of-the-art results. Currently DGRNA only has 100 million parameters, which is fewer than most RNA language models. As we continue to explore the capabilities of DGRNA with more number of pre-training RNA sequences and parameters, we expect that it will be further improved and serve as a valuable tool in the field of RNA research, contributing to our understanding of RNA’s role in various biological processes and diseases, paving the way for the development of pre-trained language models for other biomolecules such as proteins and DNA.

## Methods

### Pretraining Data of DGRNA

To construct pretraining data, we collected 1.7 billion sequences from MARS database (http://zhouyq-lab.szbl.ac.cn/download/). The MARS database integrates RNA sequences from RNAcentral, transcriptome and metagenome assemblies from MG-RAST[31], genomic sequences from the Genome Warehouse (GWH) [32]and MGnify[33], as well as NCBI’s nucleotide database (nt) [34]. As a result, MARS is 20 times larger than NCBI’s nt database and 60 times larger than RNAcentral. Due to the large volume of data, we opted not to use CD-HIT but instead chose the faster MMseqs to remove redundant data. Finally, we obtained 1.2 billion non-redundant RNA sequence[35, 36]. We keep RNA sequences longer than 3 and shorter than 2048, randomly selected to obtain a total of approximately 100 million RNA sequences for RNA language model training.

### Tokenization scheme

We employed pre-processing all non-coding RNA (ncRNA) sequences by substituting thymine (‘T’) with uracil (‘U’) as the RNA-FM. Consequently, this results in a dataset with four primary types of bases, leading to a total of 16 possible base combinations: ‘A’, ‘C’, ‘G’, ‘U’, ‘R’, ‘Y’, ‘K’, ‘M’, ‘S’, ‘W’, ‘B’, ‘D’, ‘H’, ‘V’, ‘N’, and ‘-’, “[CLS]” token and “[EOS]” token are placed at the beginning and end of the sequence, respectively[37].

### The Mamba2 Architecture in DGRNA

We build our pretraining model based on Mamba2, which has proven to be a highly effective alternative for sequence analysis tasks. Mamba is a novel SSM that offers a sub-quadratic complexity alternative to CNNs, RNNs, and Transformers, effectively modeling long-range sequential dependencies. It features a selective scanning mechanism that scales linearly with sequence length, reducing computational costs significantly compared to Transformer-based models. To enhance SSM efficiency, Mamba2 is an improved version by leveraging the theoretical SSD framework connecting SSM, attention mechanisms, and structured matrices. In DGRNA, we build upon the Mamba2 module to propose a novel bidirectional Mamba2 module, which includes forward and backward components. Each component integrates SSD elements to process sequences from left to right and right to left, respectively, followed by flash-attention and two linear layers for further processing. DGRNA consists of 12 Mamba2 layers, 1 attention heads with a total of 100 million model parameters. For a DGRNA sequence of length L, the framework generates an embedding matrix of size L×768.

Mamba2 is an advanced computational framework built on the State Space Dual (SSD) framework, designed for processing long sequence data. This framework leverages the properties of semi-separable matrices, offering an efficient way for representing and handling sequence data:

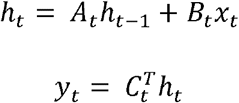

where A_t_ is the matrix controlling the temporal dynamics at timestep t.

### Bidirectional Mamba2

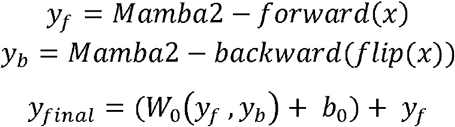

where W_0_, is the weight b_0_ is bias.

### Flash attention in DGRNA

The combination of Flash Attention with the mamba2 framework has been proven to be an effective strategy[28, 38]. Therefore, we have designed an IO-aware FlashAttention-2 layer to further reduce memory usage and computational load, thereby accelerating large-scale attention computations. This is an algorithm used to improve the efficiency of attention mechanisms. The Flash Attention layer is positioned at the end of the DGRNA model, before two linear layers, with 32 attention heads.

### DGRNA training details

We trained our DGRNA model using four RTX 3090 GPUs, each with 24 GB of memory, utilizing fairseq toolkit[39]. The training employed an inverse square root learning rate schedule with a base learning rate of 0.0001, a weight decay of 0.01, and 1,000 warm-up steps. To optimize the memory usage and speed up model training, we set the maximum input sequence length to 2048 with the batch size of 16.

During model pre-training, we utilized a self-supervised approach similar to BERT, where 15% of nucleotide tokens are randomly corrupted. Specifically, for each chosen token, 80% are replaced with a [MASK] token, 10% are substituted with a random token from the vocabulary, and the remaining 10% are left unchanged. This corruption strategy helps in training DGRNA to learn meaningful representations of nucleotide sequences.

### Downstream task datasets

#### Mean Ribosome Loading Dataset

To efficiently translate mRNAs into proteins, cells often use groups of ribosomes called polyribosomes, with the mean ribosome load (MRL) metric quantifying this process.

Usually, the translation efficiency of 5’ untranslated region (5’ UTR) is measured with MRL values. We collected a 5’ UTR dataset similar to RNA-FM, which includes two subsets: Random7600 and Human7600. These subsets consist of sequences ranging from 25 to 100 nucleotides in length. The dataset contains random 7,600 sequences for the test set and 7,600 real human 5’ UTRs. Additionally, 76,319 sequences are included in the training set[40].

#### Non-coding RNA Dataset

Approximately 5% of RNA transcripts function as mRNAs, which code for proteins, while the majority are non-coding RNAs (ncRNAs). Accurate modeling of ncRNA types is crucial for understanding their roles and the biological processes. In this study, we utilized two distinct sources of datasets: one, referred to as **non-coding_s1**, was obtained from Uni-RNA, and the other, known as **nRC**, was downloaded from RNAErnie. Among these, **non-coding_s1** was constructed as described by Cerulo et al., which involved randomly partitioning the data into three subsets: training (84%), validation (8%), and testing (8%). To mitigate the potential bias, the protocol ensured that all sequences in the validation and test sets had a normalized Hamming distance of less than 0.50 from any sequence in the training set. **nRC** comprises non-coding RNA (ncRNA) sequences selected from the Rfam database, encompassing 13 different types. The dataset includes 6,320 sequences for model training and 2,600 sequences for testing[41, 42].

#### RNA-RNA Interaction Dataset

Since most target sites are believed to be located in the 3’ UTR and most method specifically considers these regions. We assess our model’s performance using benchmark datasets: DeepMirTar, followed by the RNAErnie approach. DeepMirTar focuses on predicting interactions between microRNAs (miRNAs which are all shorter than 22 nucleotides (nts)) and mRNAs (are shorter than 53 nts). This dataset contains 13,860 positive pairs and 13,860 negative pairs[43].

#### Translation efficiency and mRNA expression dataset

The level of protein expression is influenced by both the mRNA expression level (EL) of the transcripts. To evaluate this, we utilized a benchmark dataset from UTR-LM, which incorporates three endogenous datasets for training and test: human muscle tissue (Muscle), the human prostate cancer cell line PC3 (PC3), and the human embryonic kidney 293T cell line (HEK). For sequences longer than 100 bp, portions were extracted from both the 5′ and 3′ ends to generate two separate 100 bp 5′ UTRs. For those shorter than 100 bp, the sequences were filled with repeats of the CAA motif, which is known to have no complex secondary structures. This strategy was employed to standardize the length of the sequences for subsequent analysis, while also ensuring that the added sequences do not introduce secondary structures that could influence translation efficiency[44].

#### RNA protein binding dataset

Here, we chose the PrismNet benchmark dataset for our experiments and comparisons on RNA-protein interaction prediction. PrismNet is a multi-model deep learning model used to predict the interactions between RNA-binding proteins (RBPs) and RNA. It utilizes experimental secondary structures, such as icSHAPE, to generate structural information about RNA. These data are categorized into multiple sub-datasets based on different corresponding RBPs and specific cellular contexts. Ultimately, we selected 17 RBPs, similar to RNA-FM and ERNIE-RNA, with their quantities ranging from 1,827 to 15,002. The data follows an 80:20 split ratio. To ensure consistency for input, the lengths of the different RBP datasets were standardized to 101 nucleotides[45].

#### Splicing Site Dataset

RNA splicing is crucial for gene expression, involving the removal of introns and the joining of exons. Identifying donor and acceptor splice sites is essential and is typically formulated as a binary classification task. Our model’s training dataset is obtained from Spliceator, which utilizes the G3PO+ benchmark from a wide range of eukaryotic organisms. Spliceator also provides five independent test datasets from a variety of organisms, including vertebrates (humans and zebrafish), invertebrates (fruit flies and worms), and plants (Arabidopsis). Evaluation metrics are similar to SpliceBERT and RNA-FM[46, 47].

---

**Supplementary Table 1.**
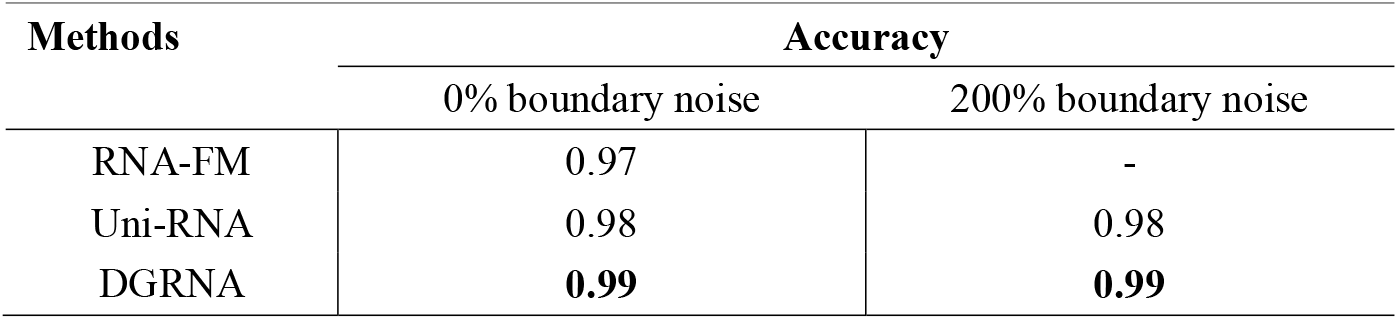
The different RNA foundation models were evaluated on two subsets of **non-coding_s1** with the metric accuracy.

| Methods | Accuracy |  |
| --- | --- | --- |
|  | 0% boundary noise | 200% boundary noise |
| RNA-FM | 0.97 | - |
| Uni-RNA | 0.98 | 0.98 |
| DGRNA | **0.99** | **0.99** |

**Supplementary Table 2.**
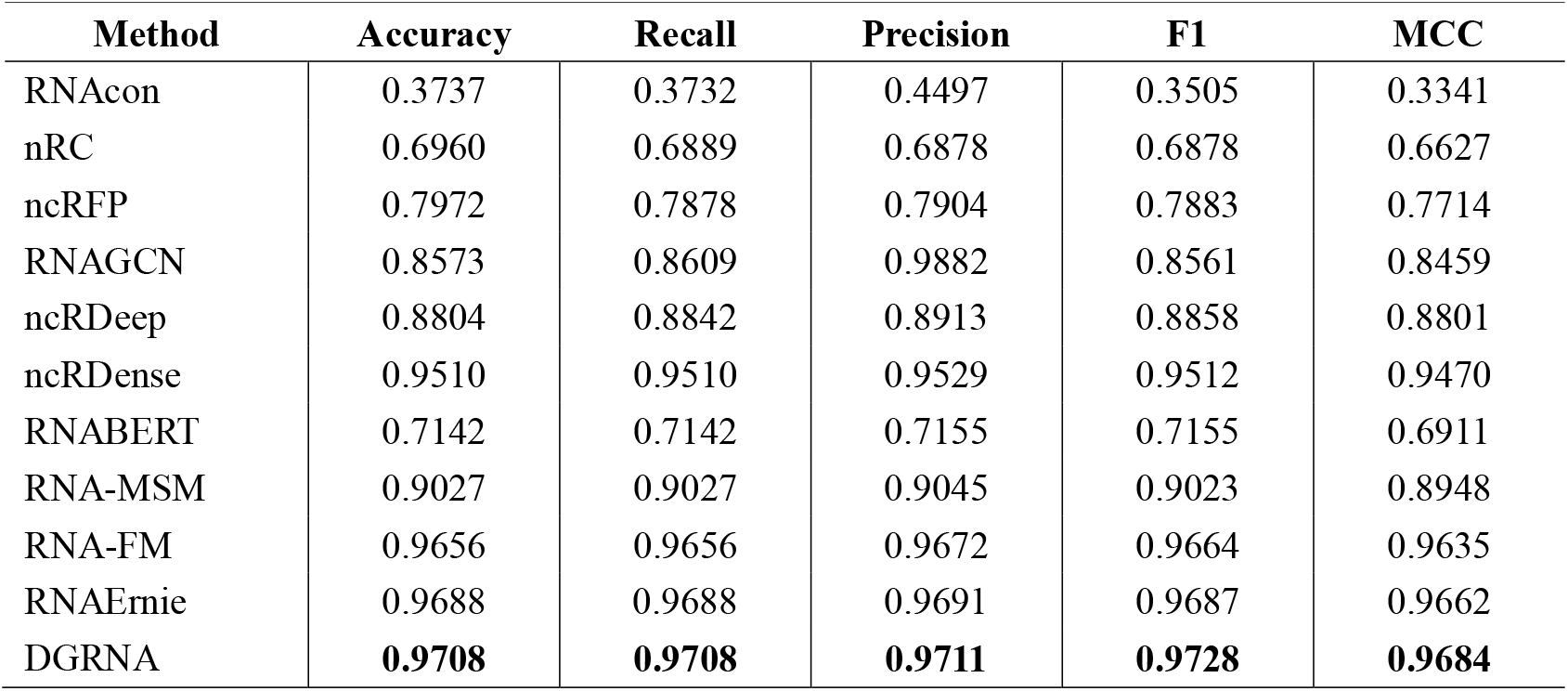
The different RNA models were evaluated on nRC dataset.

| Method | Accuracy | Recall | Precision | F1 | MCC |
| --- | --- | --- | --- | --- | --- |
| RNAcon | 0.3737 | 0.3732 | 0.4497 | 0.3505 | 0.3341 |
| nRC | 0.6960 | 0.6889 | 0.6878 | 0.6878 | 0.6627 |
| ncRFP | 0.7972 | 0.7878 | 0.7904 | 0.7883 | 0.7714 |
| RNAGCN | 0.8573 | 0.8609 | 0.9882 | 0.8561 | 0.8459 |
| ncRDeep | 0.8804 | 0.8842 | 0.8913 | 0.8858 | 0.8801 |
| ncRDense | 0.9510 | 0.9510 | 0.9529 | 0.9512 | 0.9470 |
| RNABERT | 0.7142 | 0.7142 | 0.7155 | 0.7155 | 0.6911 |
| RNA-MSM | 0.9027 | 0.9027 | 0.9045 | 0.9023 | 0.8948 |
| RNA-FM | 0.9656 | 0.9656 | 0.9672 | 0.9664 | 0.9635 |
| RNAErnie | 0.9688 | 0.9688 | 0.9691 | 0.9687 | 0.9662 |
| DGRNA | **0.9708** | **0.9708** | **0.9711** | **0.9728** | **0.9684** |

**Supplementary Table 3.** The different RNA models were evaluated on RNA-RNA interaction set.

| Method | Accuracy | Recall | Precision | F1 | AUC |
| --- | --- | --- | --- | --- | --- |
| Miranda | 0.6592 | 0.6522 | 0.6662 | 0.6591 | 0.6874 |
| RNAhybrid | 0.6988 | 0.6446 | 0.7535 | 0.6948 | 0.7585 |
| PITA | 0.4981 | 0.5872 | 0.4082 | 0.4816 | - |
| TargetScan v7.0 | 0.5801 | 0.6023 | 0.5922 | 0.5972 | 0.6725 |
| TarPmiR | 0.7446 | 0.7368 | 0.7656 | 0.7509 | 0.8021 |
| DeepMirTar | 0.9348 | 0.9235 | 0.9479 | 0.9245 | 0.9793 |
| RNABERT | 0.8375 | 0.8372 | 0.8378 | 0.8373 | 0.9160 |
| RNA-MSM | 0.8205 | 0.8203 | 0.8207 | 0.8204 | 0.9048 |
| RNA-FM | 0.9208 | 0.9208 | 0.9208 | 0.8208 | 0.9741 |
| RNAErnie | 0.9570 | 0.9576 | 0.9571 | 0.9573 | **0.9876** |
| DGRNA | **0.9654** | **0.9654** | **0.9653** | **0.9654** | 0.9859 |

**Supplementary Table 4.** The different RNA foundation models were evaluated on RBP-RNA binding datasets across 17 RBPs. The evaluation metric is AUPRCs.

| RBP name | RNA-FM | ERNIE-RNA | DGRNA |
| --- | --- | --- | --- |
| ELAVL1 | 0.950 | 0.948 | **0.955** |
| UPF1 | 0.934 | 0.943 | **0.967** |
| TIA1 | 0.929 | 0.943 | **0.953** |
| METTL14 | 0.866 | **0.912** | 0.885 |
| HNRNPC | 0.883 | 0.895 | **0.929** |
| CSTF2 | 0.867 | 0.891 | **0.931** |
| TIAL1 | 0.852 | 0.869 | **0.905** |
| U2AF65 | 0.899 | 0.921 | **0.923** |
| eIF4AIII | 0.814 | 0.853 | **0.900** |
| WTAP | 0.801 | **0.876** | 0.809 |
| HNRNPU | 0.829 | 0.849 | **0.940** |
| PTBP1 | 0.799 | 0.839 | **0.891** |
| PTBP1PTBP2 | 0.749 | 0.776 | **0.856** |
| EIF4A3 | 0.751 | 0.834 | **0.838** |
| U2AF2 | 0.705 | 0.807 | **0.863** |
| METTL3 | 0.750 | **0.832** | 0.814 |
| YTHDF2 | 0.638 | 0.725 | **0.756** |
| Mean | 0.824 | 0.865 | **0.889** |

## Reference

1. Mattick, J.S. and I.V. Makunin, Small regulatory RNAs in mammals. Hum Mol Genet, 2005. 14 Spec No 1: p. R121–32.

2. Cooper, T.A., L. Wan, and G.J.C. Dreyfuss, RNA and disease. 2009. 136(4): p. 777–793.

3. Caprara, M.G. and T.W. Nilsen, RNA: versatility in form and functio. Nat Struct Biol, 2000. 7(10): p. 831–3.

4. Morris, K.V. and J.S. Mattick, The rise of regulatory RNA. Nat Rev Genet, 2014. 15(6): p. 423–37.

5. Shang, R., et al., microRNAs in action: biogenesis, function and regulation. Nat Rev Genet, 2023. 24(12): p. 816–833.

6. Mattick, J.S. and I.V. Makunin, Non-coding RNA. Hum Mol Genet, 2006. 15 Spec No 1: p. R17–29.

7. Hombach, S. and M. Kretz, Non-coding RNAs: Classification, Biology and Functioning. Adv Exp Med Biol, 2016. 937: p. 3–17.

8. St Laurent, G., C. Wahlestedt, and P. Kapranov, The Landscape of long noncoding RNA classification. Trends Genet, 2015. 31(5): p. 239–51.

9. Cao, X., et al., Identification of RNA structures and their roles in RNA function. Nat Rev Mol Cell Biol, 2024. 25(10): p. 784–801.

10. Vervaeke, P., et al., Regulatory guidelines and preclinical tools to study the biodistribution of RNA therapeutics. Adv Drug Deliv Rev, 2022. 184: p. 114236.

11. Tants, J.N. and A. Schlundt, Advances, Applications, and Perspectives in Small-Angle X-ray Scattering of RNA. Chembiochem, 2023. 24(17): p. e202300110.

12. Jackson, R.W., C.M. Smathers, and A.R. Robart, General Strategies for RNA X-ray Crystallography. Molecules, 2023. 28(5).

13. Chen, K., et al., The Master Database of All Possible RNA Sequences and Its Integration with RNAcmap for RNA Homology Search.. 2023: p. 2023.02.01.526559.

14. Chen, K., et al., MARS and RNAcmap3: The Master Database of All Possible RNA Sequences Integrated with RNAcmap for RNA Homology Search.. Genomics Proteomics Bioinformatics, 2024. 22(1).

15. RNAcentral 2021: secondary structure integration, improved sequence search and new member databases. Nucleic Acids Res, 2021. 49(D1): p. D212–d220.

16. Kalvari, I., et al., Rfam 14: expanded coverage of metagenomic, viral and microRNA families. Nucleic Acids Res, 2021. 49(D1): p. D192–d200.

17. Devlin, J.J.a.p.a., Bert: Pre-training of deep bidirectional transformers for language understanding. 2018.

18. Vaswani, A.J.A.i.N.I.P.S., Attention is all you need. 2017.

19. Lin, T., et al., A survey of transformers. 2022. 3: p. 111–132.

20. Chen, J., et al., Interpretable RNA foundation model from unannotated data for highly accurate RNA structure and function predictions.. 2022.

21. Kalicki, C.H. and E.D. Haritaoglu, RNABERT: RNA Family Classification and Secondary Structure Prediction with BERT pretrained on RNA sequences.

22. Zhang, Y., et al., Multiple sequence alignment-based RNA language model and its application to structural inference. 2024. 52(1): p. e3–e3.

23. Yin, W., et al., ERNIE-RNA: An RNA Language Model with Structure-enhanced Representations. 2024: p. 2024.03. 17.585376.

24. Penić, R.J., et al., Rinalmo: General-purpose rna language models can generalize well on structure prediction tasks. 2024.

25. Wang, X., et al., UNI-RNA: universal pre-trained models revolutionize RNA research. 2023: p. 2023.07. 11.548588.

26. Gu, A. and T.J.a.p.a. Dao, Mamba: Linear-time sequence modeling with selective state spaces. 2023.

27. Qu, H., et al., A survey of mamba. 2024.

28. Dao, T. and A.J.a.p.a. Gu, Transformers are SSMs: Generalized models and efficient algorithms through structured state space duality.. 2024.

29. Gu, T., et al., miTAR: a hybrid deep learning-based approach for predicting miRNA target. 2021. 22: p. 1–16.

30. Wang, N., et al., Multi-purpose RNA language modelling with motif-aware pretraining and type-guided fine-tuning. 2024: p. 1–10.

31. Keegan, K.P., E.M. Glass, and F.J.M.e.g. Meyer, MG-RAST, a metagenomics service for analysis of microbial community structure and function. 2016: p. 207–233.

32. Chen, M., et al., Genome Warehouse: a public repository housing genome-scale data. 2021. 19(4): p. 584–589.

33. Richardson, L., et al., MGnify: the microbiome sequence data analysis resource in 2023. 2023. 51(D1): p. D753–D759.

34. research, N.R.C.J.N.a., Database resources of the national center for biotechnology information. 2012. 41(D1): p. D8–D20.

35. Fu, L., et al., CD-HIT: accelerated for clustering the next-generation sequencing data. 2012. 28(23): p. 3150–3152.

36. Hauser, M., M. Steinegger, and J.J.B. Söding, MMseqs software suite for fast and deep clustering and searching of large protein sequence sets. 2016. 32(9): p. 1323–1330.

37. Choi, Y., et al., High information capacity DNA-based data storage with augmented encoding characters using degenerate bases.. Sci Rep, 2019. 9(1): p. 6582.

38. Lieber, O., et al., Jamba: A hybrid transformer-mamba language model. 2024.

39. Ott, M.J.a.p.a., fairseq: A fast, extensible toolkit for sequence modeling. 2019.

40. Sample, P.J., et al., Human 5′ UTR design and variant effect prediction from a massively parallel translation assay. 2019. 37(7): p. 803–809.

41. Noviello, T.M.R., et al., Deep learning predicts short non-coding RNA functions from only raw sequence data. 2020. 16(11): p. e1008415.

42. Fiannaca, A., et al., nRC: non-coding RNA Classifier based on structural features. 2017. 10: p. 1–18.

43. Wen, M., et al., DeepMirTar: a deep-learning approach for predicting human miRNA targets. 2018. 34(22): p. 3781–3787.

44. Cao, J., et al., High-throughput 5′ UTR engineering for enhanced protein production in non-viral gene therapies. 2021. 12(1): p. 4138.

45. Xu, Y., et al., PrismNet: predicting protein–RNA interaction using in vivo RNA structural information. 2023. 51(W1): p. W468–W477.

46. Scalzitti, N., et al., Spliceator: multi-species splice site prediction using convolutional neural networks. 2021. 22: p. 1–26.

47. Chu, Y., et al., A 5′ UTR language model for decoding untranslated regions of mRNA and function predictions. 2024. 6(4): p. 449–460.

